# Charrs of the genus *Salvelinus* (Salmonidae): hybridization, phylogeny and evolution

**DOI:** 10.1101/817775

**Authors:** Alexander G. Osinov, Alexander A. Volkov, Nikolai S. Mugue

## Abstract

Evolutionary history, systematics and taxonomy of charrs of the genus *Salvelinus* and especially of the representatives of the *S. alpinus* – *S. malma* species complex remain confused that is connected with a substantial ecological and morphological flexibility of this group and with supposed ancient hybridization between some taxa. For the analysis of phylogenetic relationships and introgressive hybridization between the species of the genus *Salvelinus* including three endemic species from Lake El’gygytgyn and all main representatives of the *S. alpinus* – *S. malma* species complex, nucleotide sequences of mtDNA control region (960 bp) and two nuclear genes (ITS1 (581 bp) and RAG1 (899 bp)) were analyzed. The differences in the topologies of individual gene trees, among others reasons, were connected with incomplete lineage sorting and historical introgressive hybridization between certain taxa. Several cases of mtDNA capture by different taxa and phylogenetic groups were proposed. In particular, the following taxa participated in introgressive hybridization: northern Dolly Varden *S. m. malma*, representatives of the *S. alpinus* complex (including mainly Taranets charr *S. a. taranetzi*), southern Dolly Varden *S. m. lordi* from North America and bull trout *S. confluentus.* Main phylogenetic groups of the *S. alpinus* – *S. malma* species complex were revised. The origin and phylogenetic relationships of southern Dolly Varden from North America were not unambiguously defined. We proposed that introgressive hybridization had an important role in the evolutionary history of charrs, in particular, in the appearance of a high level of morphological, ecological and taxonomical diversity.

## INTRODUCTION

The genus *Salvelinus* is one of the most problematic (from the systematics point of view) taxon within the subfamily Salmoninae. Tentatively, all species of this genus can be subdivided into two groups. The first group includes several species whose status is not doubtful. They are as follows: brook trout *S. fontinalis* (Mitchill), lake trout *S. namaycush* (Walbaum), bull trout *S. confluentus* (Suckley), Levanidov’s charr *S. levanidovi* Chereshnev, Skopetz et Gudkov and white-spotted charr *S. leucomaenis* (Pallas). This group also includes long-finned charr *S. svetovidovi* Chereshnev et Skopetz, which has been initially separated into a distinct genus *Salvethymus* (Chereshnev and Skopets 1990). The modern ranges of the three first species are located in North America, and the other species are distributed in Eurasia. The sizes of the ranges differ substantially. The smallest ranges are known in long-finned charr, endemic species of Lake El’gygytgyn, and Levanidov’s charr with the modern range restricted by several rivers of the Sea of Okhotsk basin. In the phylogenetic tree of the genus *Salvelinus*, the species of this group have a basal position. Their relationships can be changed depending on the set of molecular markers and methods of phylogenetic analysis (e.g. Phillips and Oakley 1997; Brunner et al. 2001; Crespi and Fulton 2004; Crete-Lafreniere et al. 2012; Osinov et al. 2015; Oleinik et al. 2015). The topological position of bull trout is the most uncertain.

The second species group includes Arctic charr *S. alpinus* (L.), Dolly Varden *S. malma* (Walbaum) and relative forms and species initially referred to the S. alpinus complex (McPhail 1961; Savvaitova and Volobuev 1978; Behnke 1972, 1980). In the subsequent revision of this group, to take into account the species validity of Arctic charr and Dolly Varden, Behnke (1989) separated all taxa of the group into two sister species complexes: *S. alpinus* complex and *S. malma* complex. The representatives of the former complex have a circumpolar range with minimum penetration into the Pacific Ocean basin. The range of the latter complex is restricted to the Pacific basin, and only northern Dolly Varden is distributed (in addition to North Pacific) in the adjacent areas of the Arctic basin. According to some authors (e.g., Phillips and Oakley 1997; Osinov 2001), the two complexes can be combined into a single supercomplex *S. alpinus* – *S. malma* complex. This opinion is based on two reasons. At first, the most likely center of origin of the supercomplex is the Pacific Ocean basin, and southern Dolly Varden from Asia is the most similar to the common ancestor (Osinov and Pavlov 1998; Osinov 2001). Secondly, both the origin of southern Dolly Varden from North America and its belonging to the *S. malma complex* is questionable (Osinov et al. 2015). When the origin and phylogenetic position of southern Dolly Varden from North America will be determined, the relevance of the *S. alpinus* – *S. malma* complex concept seems not essential any more.

The number of species separated within the *S. alpinus* complex or the supercomplex, in general, substantially depends on the point of view of a particular researcher. For example, from 20 to 30 species have been described within the *S. alpinus* complex (Berg 1948; Glubokovsky et al. 1993; Chereshnev et al. 2002; Kottelat and Freyhof 2007), and more than 40 potentially valid species names of the genus *Salvelinus* including 30 representatives of the *S. alpinus* – *S. malma* complex is given in FishBase ver. 06/2018 (http://www.fishbase.org/). This species number, certainly, is not limited to take into account a great number of morphologically and ecologically distinctive charr populations including reproductively isolated sympatric forms often observed in different lakes along the entire range of the species complex. In the majority of cases, both mechanisms of the origin of these forms and levels of their reproductive isolation have not been defined. Therefore, the assignment of the species status for these forms without population genetic and phylogenetic analyses seems premature. In addition, the joining of several forms from different areas of the range into a single taxon only on the base of their morphological similarity can be erroneous. For example, ‘Boganida’ charr from Lake El’gygytgyn (Viktorovsky et al. 1981) was mistakenly combined with Boganida charr (*S. boganidae* Berg) from Taimyr described previously as a full species (Osinov et al. 2015). An assumption of Behnke (1984, 2002) on the possible common origin of Arctic charr populations of high Arctic region from the area between the Ob River in the west and Labrador in the east and their belonging to a single subspecies *S. a. erythrinus* also seems erroneous. Based on mtDNA analysis, the majority of Arctic charr populations from Siberia carry the haplotypes of Siberia group, but several populations from Chukotka and Arctic charr from Arctic regions of North America and western Greenland have the haplotypes of Arctic group (Brunner et al. 2001). If the main phylogenetic groups of Arctic charr would be referred to subspecies rank (e.g. Behnke 1980, 1984), the Siberian populations should belong to the subspecies *S. a. erythrinus*, and the populations from Chukotka, Kamchatka, western Greenland, and North America (with the exception of a part of populations from Atlantic coast (*S. a. oquassa*)) should be described as *S. a. taranetzi* (Osinov 2001; Osinov et al. 2003). The phylogenetic group of Taranets charr also includes some species with local ranges: *S. elgyticus* Viktorovsky et Glubokovsky and *S. ‘boganidae’* from L. Elgygytgyn, *S. neiva* Taranetz from Okhota R. (Osinov et al. 2015, 2018), as well as charr populations from Alaska designated previously as a ‘Bristol Bay-Gulf of Alaska’ form (Behnke 1980, 2002; Osinov 2001).

Cyclic Pleistocene glacial periods had a huge impact on the biota of northern ecosystems (Hewitt 2004) leading to substantial reconstruction, fragmentation, and disposition of the native ranges of many species. Some populations were subjected to extinction, and others were displaced into large or cryptic glacial refugia. Postglacial distribution of many animal and plant species (glacial lineages) from these refugia led to secondary contacts and hybridization including the formation of hybrid zones (Hewitt 1999, 2000, 2004). Many ichthyologists suppose that the origin of a substantial part of present biodiversity within the genus *Salvelinus* is connected with the glacial period events, but a role of historical hybridization has been assessed differently (Taranets 1936; McPhail 1961; Behnke 1972, 1980, 1984, 2002; Savvaitova 1989; Haas and McPhail 1991; Osinov 2001; Power 2002; Klemetsen 2010; Taylor 2016).

Numerous hybridization events between the forms and species of charrs have been described (e.g., Behnke 1980, 2002; Taylor 2004, 2016; Reist and Sawatzky 2010;). In some cases, a capture of ‘alien’ mtDNA occurred in distinct populations (or local groups) due to introgressive hybridization. These cases are described for different species (Wilson and Hebert 1993; Wilson and Bernatchez 1998; Bernatchez et al. 1995) and different phylogenetic groups of the *S. alpinus* – *S. malma* complex (Osinov et al. 2017, 2018; Esin et al. 2017). In various representatives of the *S. alpinus* – *S. malma* complex, ‘alien’ mtDNA, most often, belongs to the haplogroup of northern Dolly Varden (Osinov and Mugue 2008; Taylor et al. 2008; Alekseyev et al. 2009; Moore et al. 2015; Osinov et al. 2017, 2018; Esin et al. 2017). The introgression of northern Dolly Varden mtDNA, most likely, occurred during huge postglacial expansion of this form from a glacial refugium and its secondary contacts and hybridization with different representatives of the *S. alpinus* – *S. malma* complex (Osinov et al. 2015, 2018). Owing to introgressive hybridization between different charr taxa, the identification of several existing populations and forms is difficult, erroneous phylogenetic and taxonomic interpretations are possible (see examples: Osinov et al. 2015, 2018) and phylogenetic analysis becomes complicated.

The goal of this study is the analysis of phylogenetic relationships and historical hybridization events between the forms, species and phylogenetic groups of charrs of the genus *Salvelinus* and especially of the *S. alpinus* – *S. malma* complex. We will describe the most probable cases of introgressive hybridization, which occurred both during the postglacial period and in earlier ages, and its effect on the phylogenetic position of different taxa will be assessed. For this purpose, we chose three “good” markers from *Salvelinus* phylogeographic and phylogenetic studies: the control region of mtDNA and two nuclear DNA fragments (RAG1 and ITS1). These markers have a good phylogenetic signal even though the signals from different genes are not always the same for some taxa (or certain populations) (Phillips et al. 1995; Phillips et al. 1999; Brunner et al. 2001; Shedko et al. 2012; Osinov et al. 2018). To differentiate a possible effect of incomplete lineage sorting and introgressive hybridization, we used a parsimony approach (Yu et al. 2013) extending the ‘minimizing deep coalescence’ criterion (Maddison 1997; Than and Nakhleh 2009) for species tree and networks inference implemented in the PHYLONET package (Than et al. 2008).

## MATERIAL AND METHODS

### Sampling

Tissue samples (fin fragments or a part of muscles) of the individuals of different taxa of the genus *Salvelinus* including all main representatives of the *S. alpinus* – *S. malma* complex were collected within the entire range of the genus from 1985 to 2015 (Table 1, Fig. 1). All samples were obtained from natural populations of different species and phylogenetic groups, but the boundaries of the *S. alpinus* – *S. malma* complex ranges had not been determined precisely (Online Resource 1). The tissue samples of *S. fontinalis* and *S. namaycush* were collected from the broodstocks.

**Figure 1.**
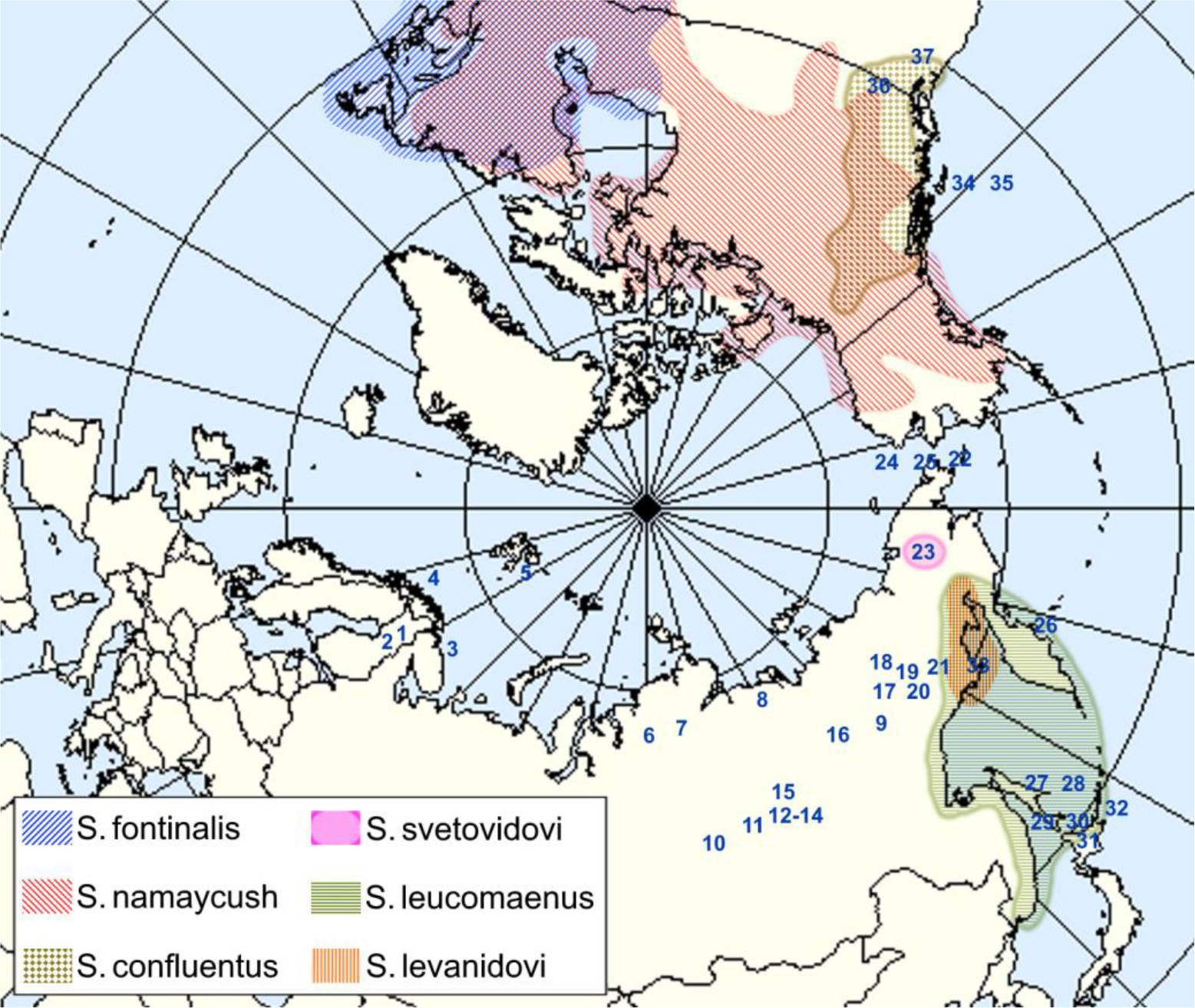
Sampling locations of *Salvilinus* taxa (according to the data from Table 1) and current ranges of *S. fontinalis*, *S. namaycush, S. confluentus*, *S. svetovidovi*, *S. leucomaenus* and *S. levanidovi*. 1, Luomusjarvi Lake; 2, Kanes Laddu Lake; 3, Vyashenskoe Lake; 4, Fjellfrosvatnet Lake; 5, Spitsbergen; 6, Lama Lake; 7, Ayan Lake; 8, Baganytta-Kyuel Lake; 9, Dyampa-Kyuel Lake; 10, Svetlinskoe Lake; 11, Bol’shoi Namarakit Lake; 12, Leprindokan Lake; 13, Bol’shoe Leprindo Lake; 14, Davatchan Lake; 15, Usu Lake; 16, Ulakhan-Silyan-Kyuel Lake; 17, Kobuyma-2 Lake; 18, Urasalakh-Kyuel Lake; 19, Kengre Lake; 20, Malyk Lake; 21, Juliette Lake; 22, Naivak Lake; 23, El’gygytgyn Lake; 24, Netteveem River; 25, Enurmino lagoon; 26, Kamchatka River; 27, Tym’ River; 28, Longari River; 29, Rogatka River; 30, Solov’evka River; 31, Shikaribetsu Lake; 32, Unnamed Brook; 33, Yama River; 34, Cache Creek; 35, Mountain Creek; 36, Hill Creek; 37, Yakima River.

### DNA isolation, PCR and sequencing

Total DNA was extracted from each specimen’s tissue, using Wizard SV Genomic DNA Purification System. Sequences of mtDNA control region (CR mtDNA) were obtained by amplification, using HN20 – Tpro2 primer pair described by Brunner et al. (2001). For RAG1 exon 2 dataset, we used the primer pair RAG1F (Quenouille et al. 2004) and RAG-RV1 (Šlechtová et al. 2004). ITS1 sequences dataset was obtained by using KP2 – 5.8S primer pair (Phillips et al. 1995). Standard PCR was carried out in 15 uL total volume, containing 50-100 ng of DNA template, 1.5 uL of 10X PCR buffer (670 mM Tris-HCl pH 8.8, 166 mM (NH_4_)_2_SO_4_, 0.1% Tween-20), 0.75 uL MgCL_2_, 1 uL of dNTP mix (2.5 mM of each), 0.5 uL of each primer (0.01 mM), and 0.5 uL of SmarTaq Polymerase (Dialat Ltd., Russia). The PCR was performed using the following cycling conditions: 2 min at 96°C; 35 cycles of 10 sec at 96°C, 30 sec at 50°C (for CR) or 58°C (for RAG1 ex.2) or 52°C (for ITS1), and 1 min at 72°C; 5 min at 72°C. After PCR was accomplished, the products were visualized in 2% agarose gel and, then threatened with 5 U of Exonuclease I at 37°C for 15 min, followed by 10 min at 80°C. Sequencing was performed in both forward and reverse directions using BigDye v.1.1 Cycle Sequencing Kit (Applied Biosystems, Inc.) on the AB3500 capillary sequencer. The primers for sequencing were as follows: for CR mtDNA – HN20, Tpro2 and primer CHAR3 (Alekseyev et al. 2009); for RAG1 exon 2 – RAG1F and RAG-RV1; and for ITS1 – KP2 and 5.8S.

### Data Analyses

Each sequences dataset was edited and aligned using Geneious v.6.4 (https://www.geneious.com) and Clustal X (Thompson et al. 1997) software. Heterozygous positions were coded using IUB ambiguity codes. Sequence data were deposited in GenBank (http://www.ncbi.nlm.nih.gov), corresponding to accession numbers: MK026466– MK026552.

We applied the Akaike information criterion (AIC) in the jModeltest program (Posada, 2008) for each sequence dataset to select the optimal nucleotide substitution model. The program PartitionFinder ver.1.1.0 (Lanfer et al. 2012) we used for choosing an appropriate partitioning scheme for two concatenated nuclear dataset (ITS1+RAG1). To select the optimal substitution model for the maximum likelihood (ML) reconstruction in TREEFINDER ver. March 2011 (Jobb 2011) we used the “linked branch lengths” option and AIC criterion, and for Bayesian inference (BI) in BEAST program – the “unlinked branch lengths” option and BIC criterion.

### Phylogenetic tree reconstruction

In the case of nuclear sequence data, we performed phylogeny reconstruction based on two methods of accounting for heterozygous sequences: the use of ambiguity IUB-IUPAC characters and unphased haplotype data due to all the charr specimens had no more than one heterozygous site per individual sequence in RAG1 and ITS1 datasets. To construct different types of phylogenetic trees, we used three distinct haplotype datasets. The first entire dataset included sequences of 97 charr specimens (Table 1). The second (reduced) dataset was comprised of sequences of 63 charr specimens with haplotypes, differing, at least, in one sequence dataset, and the individuals with identical haplotypes simultaneously in all three sequence datasets were excluded. Alternatively, reduced dataset consisted of 57 specimens sequences and represented the previous dataset with excluded specimens of Arctic charr and southern Dolly Varden from Asia with Bering (BER) mtDNA haplotype lineage of northern Dolly Varden. The haplotype datasets were combined with DNASP v.5 (Librado and Rosas 2009). We used *Oncorhynchus masou* as an outgroup for the tree reconstruction (GenBank accession numbers: DQ864464, GQ871486, AF170536).

An unweighted maximum-parsimony (MP) analysis was performed in PAUP v.4.0b10 (Swofford, 2002) and using haplotype data (from 63 and 64 (+*O. masou*), 57 and 58 (+*O. masou*) individuals) and the following options: nreps=10, addseq=random, MaxTrees= 1000, swap=tbr. An incongruence length difference (ILD) test (Farris et al. 1995) with 1000 replicates we used to check for significant discordances among pairs of two and all three gene trees. ML analysis for individual genes was performed in PAUP. The appropriate substitution models from jModeltest were used for heuristic search with TBR algorithm for the best ML tree. ML tree for concatenated (RAG1+ ITS1) sequences data with the best model from PartitionFinder was estimated with TREEFINDER. The best partition substitution models were: for ITS1 (1st pos.-TrN, 2nd pos.-TrN+I, 3rd pos.-HKY+I) and GTR+G+I for RAG1 (all positions). To test the node stability in ML and MP trees, a non-parametric bootstrap analysis (Felsenstein, 1985) with 500 and 1000 pseudo-replicates, respectively, was used.

Bayesian phylogenetic analysis was performed with BEAST v.1.8.4 (Drummond and Rambaut 2007). For concatenated (ITS1+ RAG1) haplotype data (from 64 individuals) we used the best substitution model (GTR+G+I) obtained in PartitionFinder with Yule tree prior and “uncorrelated lognormal relaxed clock” model. For species tree estimation we used *BEAST (Heled and Drummond, 2010) with two sets of multilocus data. The first set included RAG1 and ITS1 sequences of 97 specimens of the charrs (Table 1) grouped into 13 taxa (comprising the species and subspecies). The specimens of two phylogenetic groups (*S. a. alpinus* and *S. a. erythrynus*) were joined into a single taxon *S. alpinus* (Eurasia) due to no sequence difference of these genes. We used “Birth and Death” model (Gernhard 2008) as a tree prior, “Piecewise constant” as population size model and “Random local clock” model. For RAG1 and ITS1 datasets, the most appropriate substitution models were HKY and GTR+G respectively. The second dataset included CR, RAG1 and ITS1 haplotypes of 57 individuals grouped into 15 taxa. The southern Dolly Varden of North America with two mtDNA haplogroups (Bering, BER and East Pacific, EP) were conditionally divided into two taxa. We used the “Birth and Death” model as a tree prior, “Piecewise constant” as population size model and “uncorrelated relaxed clock” model. Each of three sequence datasets had GTR+G substitution model. Every Bayesian analysis included two independent runs with 120 million MCMC generations each with first 12 million MCMC steps discarded as burn-in. The results were checked for adequate MCMC mixing and sufficient sampling of priors using Tracer v.1.6 (Rambaut and Drummond 2007). We used TreeAnnotator v. 1.6 to combine and to summarize results from two independent runs and FigTree v.1.4.2 (http://tree.bio.ed.ac.uk/software/figtree/) for viewing the trees.

### Inferring introgressive hybridization

At this step, we employed two approaches that we implemented in two stages. The first stage, according to the mito-nuclear and mito-morphological discordance (Osinov et al. 2017, 2018), we excluded from the analysis those individuals of Arctic char and southern Dolly Varden from Asia which had mtDNA haplotypes of northern Dolly Varden (BER) as a result of introgressive hybridization (Table 1). Thus, we used on the second stage the reduced dataset of 57 specimens. Initially, in PAUP we obtained MP trees based on every gene dataset and stored them in Newick format. The branch lengths were ignored. After that, all the specimens relating to the same taxon were joined together. We obtained 15 taxa, and the individuals of southern Dolly Varden of North America which had mtDNA haplotypes BER and EP were split into two distinct taxa. We used PhyloNet (Than et al. 2008) to provide a parsimonious inference (Yu et al. 2013) with “Minimize Deep Coalescence” criterion to find the species tree (Infer_ST_MDC option) and to infer phylogenetic networks with 1 to 3 reticulation events (InferNetwork_MP option). Every set of reticulations events was accomplished with five independent runs. MDC-based species tree topology and all phylogenetic networks, including Rich Newick format indicating the reticulation nodes (#H), inheritance probabilities (for species contribution toward hybridization) and the number of extra lineages, are provided (Online Resource 2,3). We used Dendroscope 3.2.5 (Huson and Scornavacca 2012) to draw MDC-based species tree and phylogenetic networks.

## RESULTS

### Individual gene trees and their main topological differences

A 960-bp fragment of the mtDNA control region (CR mtDNA) had 157 variable characters including 58 parsimony-informative characters. In 97 individuals of the genus *Salvelinus*, 54 haplotypes were revealed (Fig. 2). In the charrs of the *S. alpinus* – *S. malma* complex, six haplogroups were registered (Fig. 2), and these haplogroups corresponded with six phylogenetic groups of charrs. The designations of five haplogroups were suggested before (Brunner et al. 2001), and one of them (Acadia) was absent in our study. A haplogroup of southern Dolly Varden from Asia was separated by Shedko et al. (2007) and designated as Okhotskaya. This name was suggested previously for the group including the haplotypes of northern Dolly Varden, neiva (*S. neiva*) and lacustrine charrs of the Sea of Okhotsk basin (Radchenko 2005). To avoid confusion in the description, we designate a haplogroup of southern Dolly Varden from Asia as West Pacific. The name East Pacific is used for the clade composed of the haplotypes, which are present in bull trout and several populations of southern Dolly Varden from North America. Both last designations were suggested by Oleinik et al. (2015).

**Figure 2.**
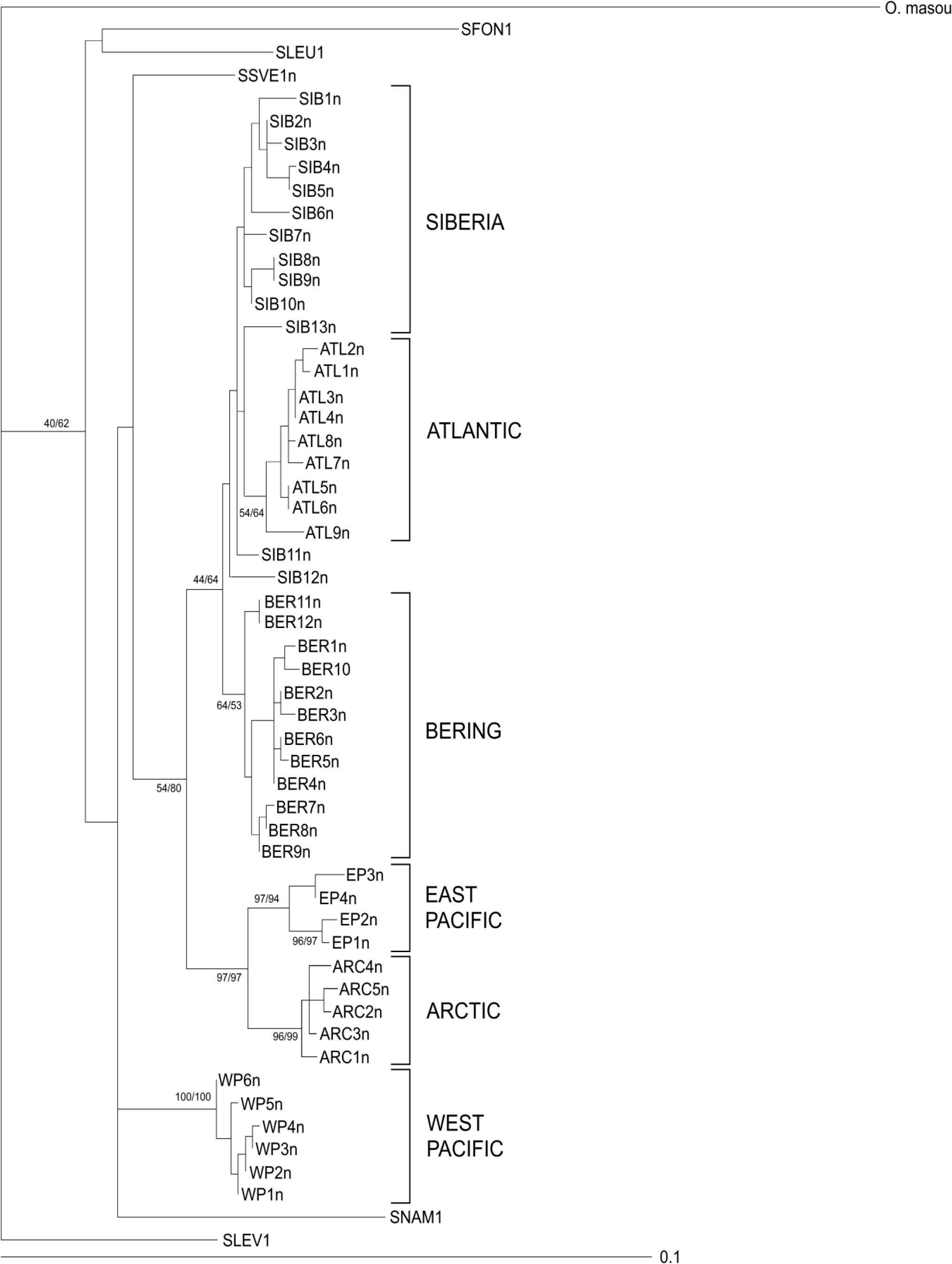
ML tree for *Salvelinus* species based on CR haplotype data and GTR+G+I substitutional model. Bootstrap values for ML/MP trees are shown on the branches. Haplotype designations see in Table 1.

The ML and MP trees of the control region had similar topologies and similar bootstrap values for the majority of clades (Fig. 2). We note the following features of the tree topologies to analyze phylogeny of the genus *Salvelinus.* The *S. alpinus* complex and *S. malma* complex are not monophyletic groups because they do not form two corresponding clades. Three clades of Dolly Varden are located at different parts of the tree. For example, the clade of the haplotypes of northern Dolly Varden (Bering group) is a sister group to the clade of the haplotypes of Arctic charr from Europe and Siberia (Atlantic and Siberia groups). The *S. alpinus* – *S. malma* supercomplex is also not monophyletic: *S. svetovidovi* and *S. confluentus* are located inside of its clade. The clade East Pacific is characterized by good bootstrap support (> 95%) and represents a sister group to a clade of Taranets charr (Arctic group). Southern Dolly Varden from Asia (West Pacific group) and *S. namaycush* are located at the base of the *S. alpinus* – *S. malma* complex clade. Levanidov’s charr is located at the base of the *Salvelinus* tree, but its separation from other taxa is not very reliable (40/62%).

A 581-bp internal ribosomal spacer ITS1 had 87 variable characters including 49 parsimony-informative characters, and 16 haplotypes were revealed in 97 charr individuals (Table 1, Fig. 3a). The ML and MP phylogenetic trees had a similar topology and following essential features. The (*S. fontinalis*, *S. namaycush*) clade is located at the base of the tree, but the bootstrap support is poor (< 50%). The (*S. levanidovi, S. leucomaenis*) clade is separated comparatively reliably (bootstraps for the ML and MP trees are 78 and 90%, respectively), and the sister taxon to this clade is *S. confluentus* (95/87%). *S. svetovidovi* is the sister taxon to the *S. alpinus* – *S. malma* complex clade. Northern and southern (Asian) forms of Dolly Varden are reliably (92/92%) combined into a monophyletic clade. A haplotype of southern Dolly Varden from North America (AC6) is located inside the clade including the haplotypes of Arctic charr and Taranats charr (*S*. *alpinus* complex). Two species from Lake Elgygytgyn share the haplotype AC4.

**Figure 3.**
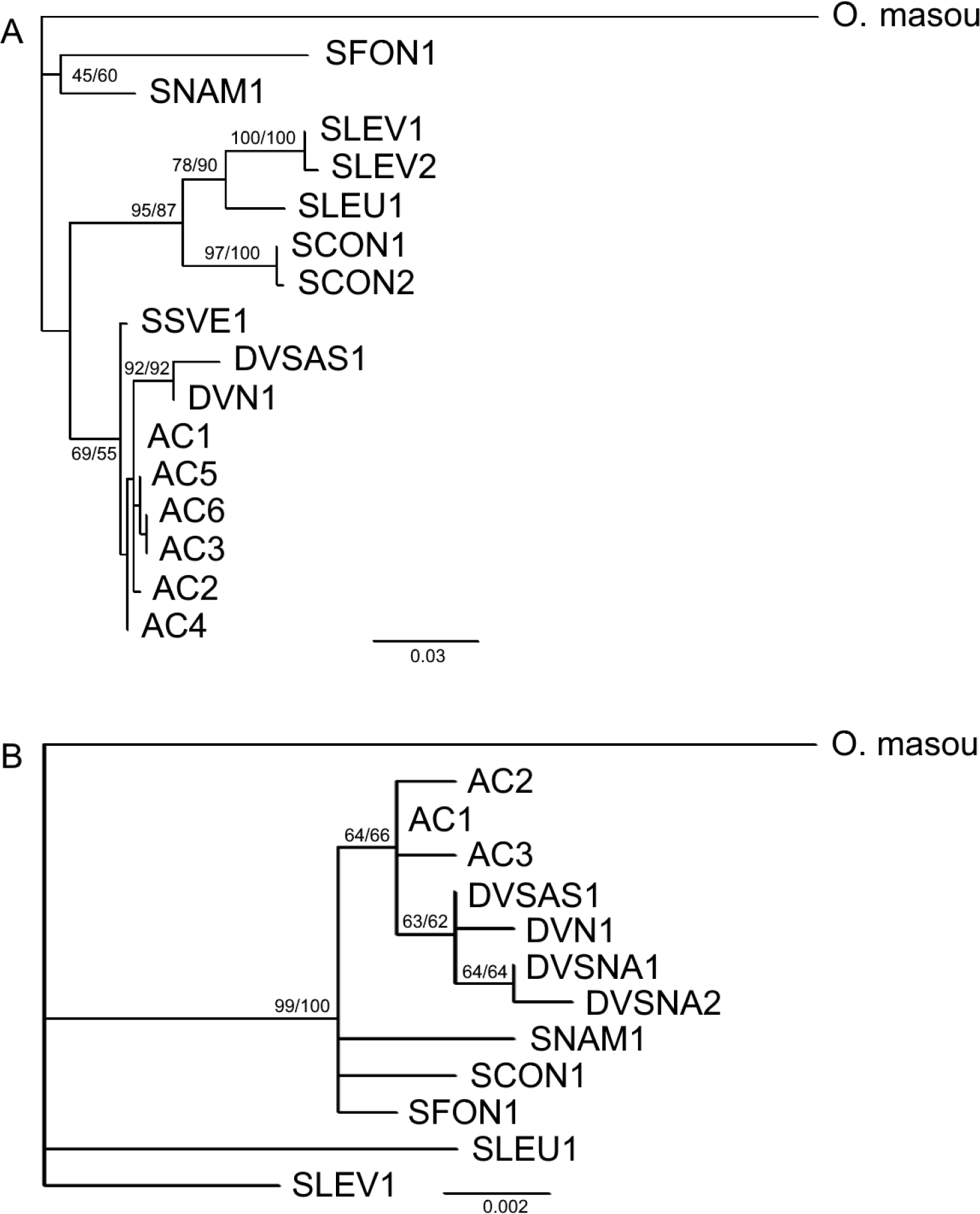
ML trees for *Salvelinus* species based on (A) ITS1 haplotypes and GTR+G+I substitutional model and (B) RAG1 haplotypes and HKY substitutional model. Bootstrap values for ML/MP trees are shown on the branches. Haplotype designations see in Table 1.

An 899-bp fragment of exon 2 of RAG1 gene had 28 variable characters including 10 parsimony-informative characters, and 12 haplotypes were revealed in the charrs of the genus *Salvelinus* (Table 1, Fig. 3b). The ML and MP tree topologies had the following essential features. *S. levanidovi* and *S. leucomaenis* are located near the root, and all other species of the genus *Salvelinus* form a clade with high bootstrap support (99/100%). Inside of this clade, a large clade including all representatives of the *S. alpinus* – *S. malma* complex is separated (bootstraps 64/66%). Long-finned charr has the AC1 haplotype, which is also revealed in sympatric *S. elgyticus*, as well as in many other representatives of the *S*. *alpinus* complex (Table. 1). All forms of Dolly Varden form a clade (*S. malma* complex) with moderate support (63/62%).

### Results of ILD test for 64 and 58 set of individuals

The results of the ILD test indicated the significant conflict among three genes (64: p = 0.0014, 58: p = 0.0015). For two genes comparison, the significant differences were registered between CR and ITS (64: p = 0.0012, 58: p = 0.0046), between RAG1 and ITS1 (64: p=0.015, 58: p=0.005), and the differences were not significant between CR and RAG1 (64: p = 0.267, 58: p = 0.492).

### Concatenated RAG1 and ITS1 data trees and species trees

The ML and MP trees constructed in PAUP had a similar topology and bootstrap support of the main clades (Fig. 4a). The species *S. fontinalis* and *S. namaycush* are located near the root. The clade ((*S. levanidovi, S. leucomaenis*), *S. confluentus*) is reliably (95/80 %) separated. *S. svetovidovi* is a sister group to the *S. alpinus* – *S. malma* complex, and *S. elgyticus* and *S. boganidae* are located at the base of this species complex clade. Southern form of Dolly Varden from Asia joins with a northern form of Dolly Varden (95/96%), and southern Dolly Varden from North America is located inside of the Taranets charr clade. The ML trees from PAUP (P) with GTR+G+I substitution model (Fig. 4a) and ML tree (not presented) from TREEFINDER (T) constructed with different substitution models for four partitions had the following topological differences. In the ML(T) tree, three charr species from Lake El’gygytgyn produce a clade *S. svetovidovi* (*S. elgyticus, S. boganidae*), which is inside of the *S. alpinus* – *S. malma* complex clade (with 97% bootstrap support). The clade including southern Dolly Varden from Asia and northern Dolly Varden (97%) is located near the base of the *S. alpinus* – *S. malma* complex clade. These two features of the ML(T) tree topology can be found in the Bayesian tree constructed in BEAST using the GTR+G+I model (Fig. 4B). One of the main differences between the topologies of the ML trees and the tree from BEAST for concatenated data is connected with the position of two basal clades (*S. fontinalis*, *S. namaycush*) and ((*S. levanidovi, S. leucomaenis*), *S. confluentus*). The latter clade has good support in all trees (Fig. 4a,b).

**Figure 4.**
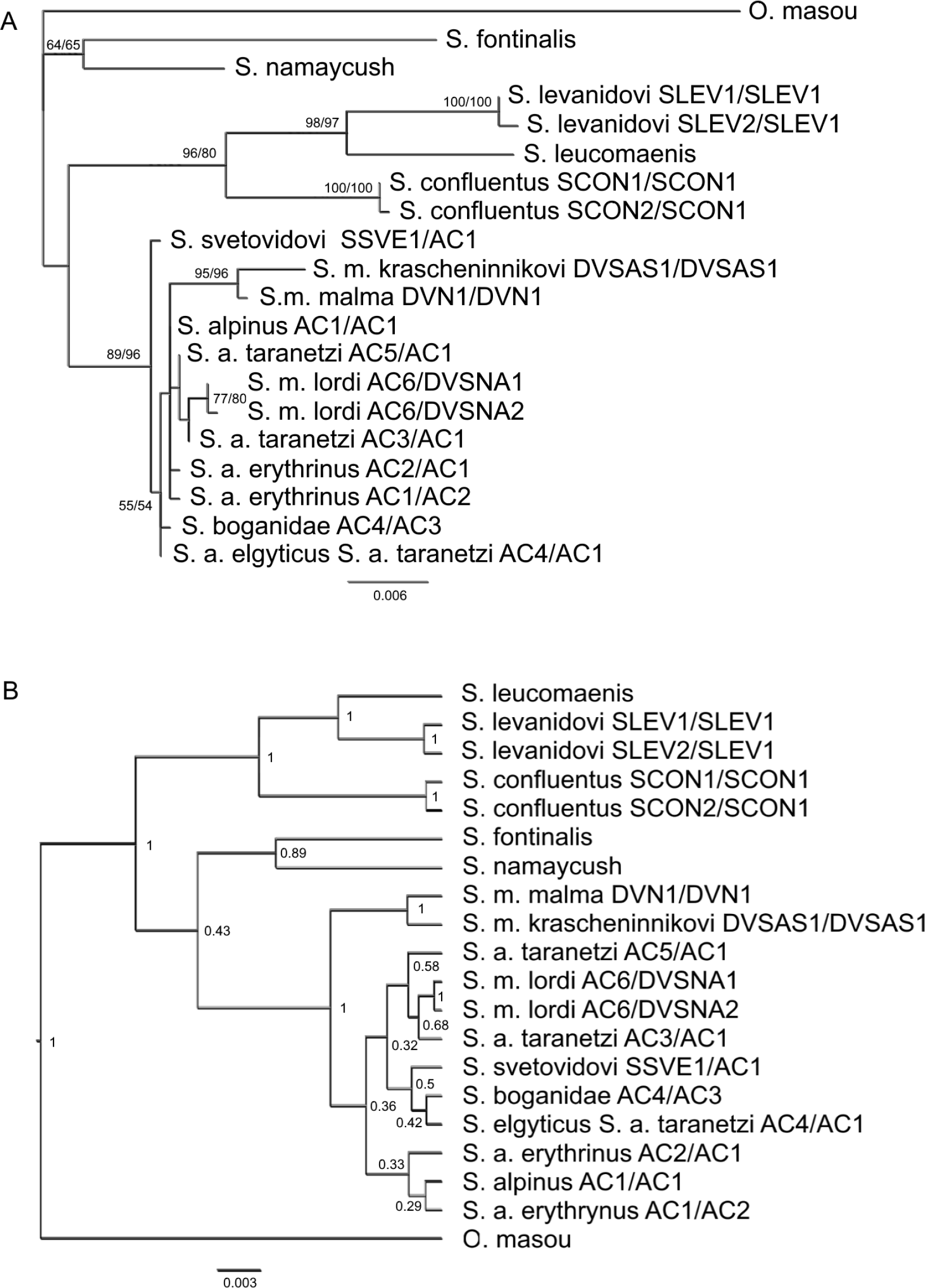
Phylogenetic relationships among *Salvelinus* species inferred from concatenated ITS1 and RAG1 haplotype data: (A) ML tree based on GTR+G+I substitution model from PAUP analysis; (B) Bayesian maximum clade credibility tree based on GTR+G+I substitution model from BEAST analysis. Numbers near nodes indicate BS or PP values. ITS1/RAG1 haplotypes are given to the right of the taxa names.

In the species tree from *BEAST analysis, based on RAG1 and ITS1 genes, the *S. alpinus* – *S. malma* complex clade has good support (0.98– posterior probability, PP). *S. svetovidovi* is located inside of this clade and represents a sister taxon to the *S. alpinus* complex clade. The latter clade includes two other charr species from Lake El’gygytgyn, Arctic charr of Eurasia and Taranets charr (Fig. 5). The support of this clade (PP = 0.21), as well as the support of many other clades of the tree, is poor. *S. confluentus* is a sister taxon to the clade (*S. levanidovi, S. leucomaenis*). In the species tree based on three genes, many clades are also characterized by low values of posterior probability (Fig. 6a). *S. svetovidovi* is a sister group to the *S. alpinus* – *S. malma* complex group, and this clade is reliably separated (PP = 0.98). Southern Dolly Varden from North America with CR haplotypes EP joins with *S. confluentus* (PP = 0.98), and the clade with CR haplotypes BER joins with *S. m. malma* (PP = 0.69). All forms of Arctic charr including two species from Lake El’gygytgyn are consolidated into a single clade (*S. alpinus* complex) with poor support (PP = 0.25).

**Figure 5.**
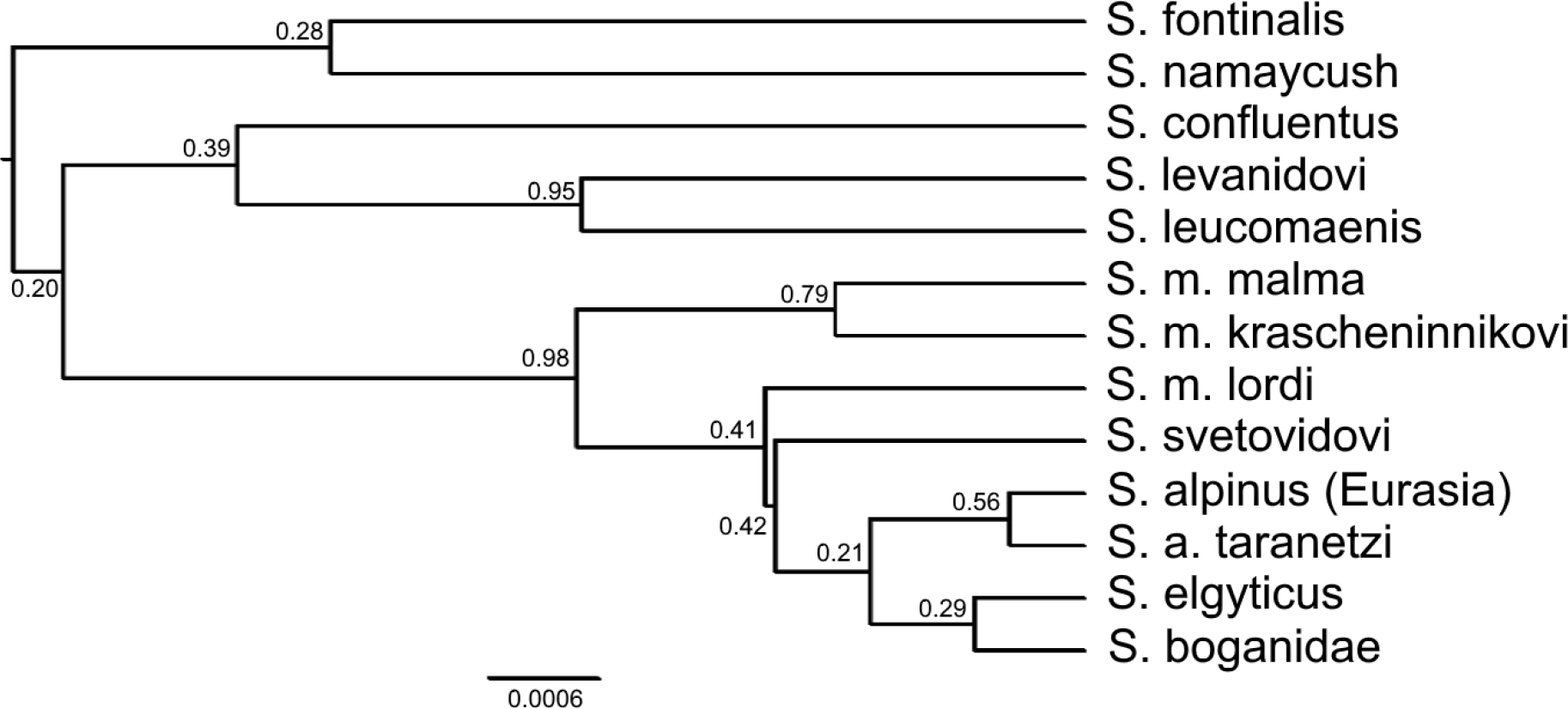
Species tree for *Salvelinus* based on two nuclear genes from *BEAST analysis. Numbers near the nodes indicate posterior probability values.

**Figure 6.**
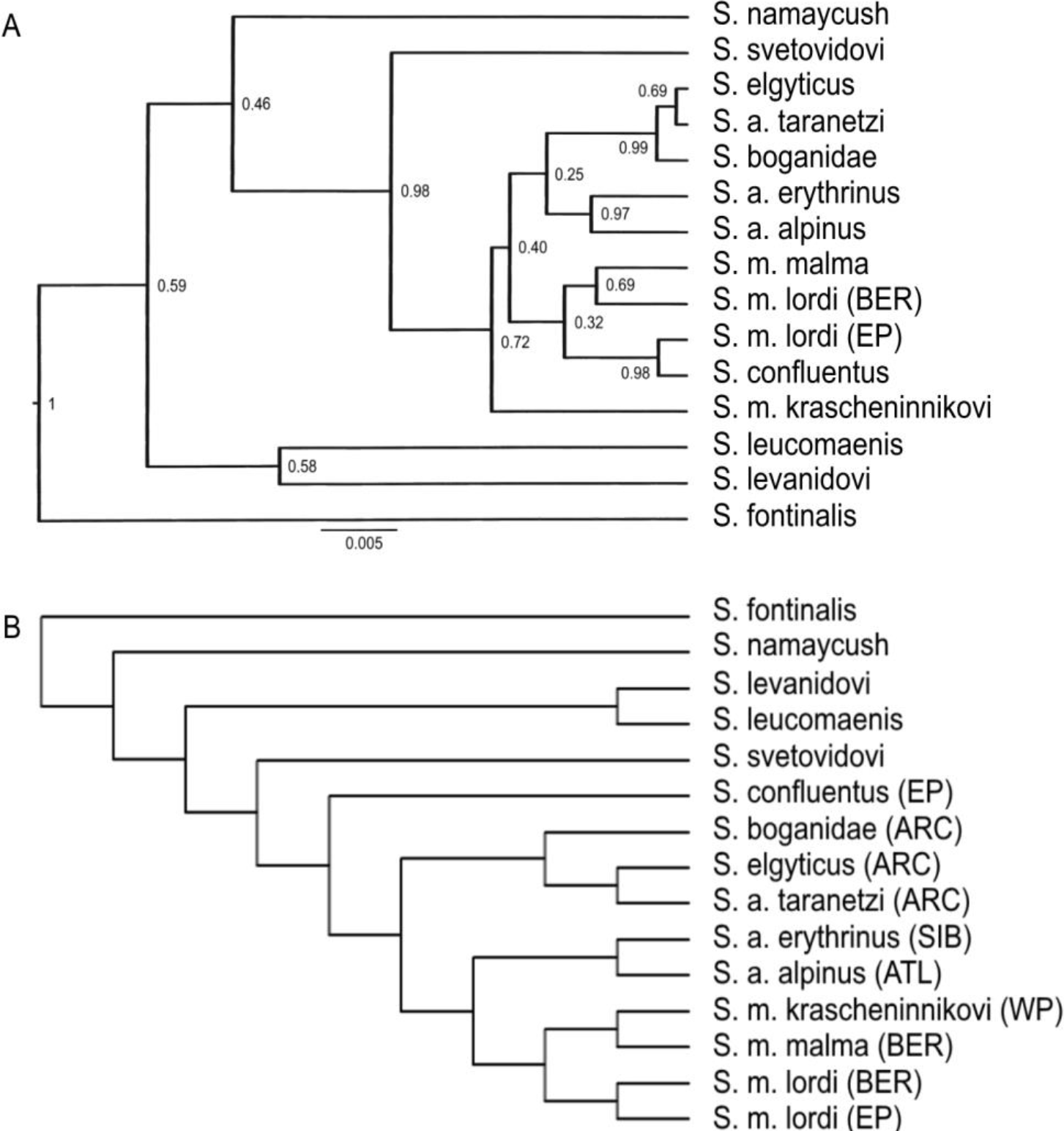
Species trees for *Salvelinus* based on three genes from (A) *BEAST and (B) Phylonet analyses. Numbers near the nodes indicate posterior probabilities. CR haplogroup is indicated in the parentheses.

### Hybridization events, species tree and phylogenetic networks from PHYLONET

The species tree inferred with MDC criterion includes 22 extra lineages (NEL) (Fig. 6b, Online Resource 2). *S. fontinalis* is located at the base of the tree followed by *S. namaycush*. *S. svetovidovi* is located between the clade (*S. levanidovi, S. leucomaenis*) and *S. confluentus*; and the latter species is a sister taxon to the *S. alpinus* – *S. malma* complex. All forms of Dolly Varden produce a single clade (*S. malma* complex) with a sister relation to the clade of Arctic charr from Europe and Siberia.

Among the phylogenetic networks obtained in five runs with one reticulation node, two networks are identical (Online Resource 2 (Fig. 2: B2, B3)). They indicate that southern Dolly Varden from North America had gene exchange with the charrs of the Taranets group on the one hand and with northern Dolly Varden on the other hand. Two other networks support historical hybridization between these taxa (Fig. 7a, Online Resource 2 (Fig. 2: B1, B4). The differences between three networks are connected with the position of *S. confluentus* and with the particular taxon of the Taranets group (a common ancestor or one of the present species) participated in hybridization with southern Dolly Varden from North America. According to a network (Online Resource 2 (Fig. 2 B5)), hybridization between *S. m. lordi* (BER) and *S. confluentus* occurred in the past. The total number of extra lineages (NEL) in five networks is 17 or 18.

**Figure 7.**
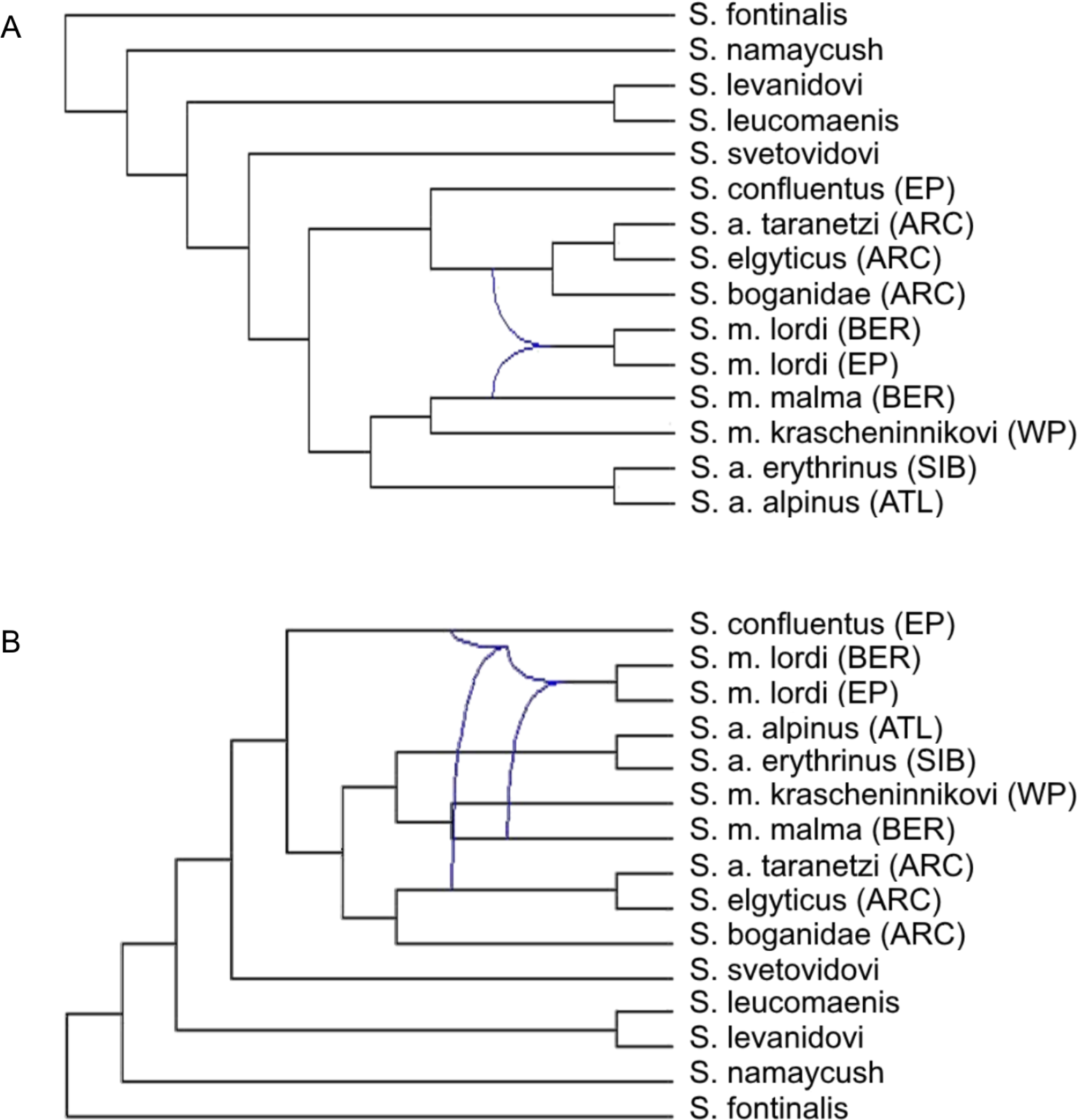
Phylogenetic networks with one (A) and two (B) reticulation nodes. CR haplogroup is indicated in the parentheses.

The network (NEL = 15) with two reticulation nodes indicates that four taxa participated in two hybridization events (Fig. 7b, Online Resource 2 (Fig. 2C1)). *S. confluentus* is added to three most likely taxa revealed in the networks with a single reticulation node. This topology is supported by two other networks (Online Resource 2 (Fig. 2 C2, C3)). One network indicates that southern Dolly Varden from North America participated in introgressive hybridization with the common ancestor of the Taranets group and common ancestor of northern and southern Dolly Varden from Asia. In addition, the common ancestor of northern and southern Dolly Varden from Asia participated in hybridization with *S. svetovidovi* and with the common ancestor of Arctic charr from Europe and Siberia (Online Resource 2 (Fig. 2C4)). A single reticulation node is seen in the network of the fifth run (Fig. S2: C5). It is supposed that southern Dolly Varden from North America went through hybridization with northern Dolly Varden and bull trout. The main difference between the topologies of various networks is connected with the position of bull trout: it represents a sister group to entire *S. alpinus* – *S. malma* complex, to only *S. alpinus* complex or only Taranets group. The total number of extra lineages in five networks varied from 15 to 18 (Online Resource 2 (Fig.2 C1-5)).

In the networks with three reticulation nodes, an unstable position of *S. confluentus* and several taxa from the *S. alpinus* – *S. malma* complex is observed (Online Resource 3 (Fig. 3)). In a network (NEL = 13), the clade of southern Dolly Varden from North America is separated into two taxa, which are located in the different parts of the network (and participated in hybridization with various taxa) (Online Resource 3 (Fig. 3.1). *S. m. lordi* (EP) participated in hybridization with *S. confluentus* and *S. m. lordi* (BER) entered into hybridization with *S. a. taranetzi* and *S. m. krascheninnikovi*. In two networks, *S. svetovidovi* participated in hybridization with (1) common ancestor of the Taranets group (Online Resource 3 (Fig. 3.4) and (2) Arctic charr from Siberia (Online Resource 3 (Fig. 3.3). In different networks, *S. confluentus* hybridized with *S. m. lordi* (Online Resource 3 (Fig. 3:1,2,4)) or with a representative of the Taranets group (Online Resource 3 (Fig. 3.3). According to a network, Arctic charr from Eurasia is not a sister group to the clade of northern and southern Dolly Varden from Asia, and only gene exchange took place (Online Resource 3 (Fig. 3.5)). The total number of extra lineages in five networks varied from 13 to 17 (Online Resource 3 (Fig. 3)).

## DISCUSSION

### Phylogenetic relationships of charrs of the genus Salvelinus and main representatives of the S. alpinus – S. malma species complex

To analyze phylogeny of charrs of the genus *Salvelinus*, different databases (morphological, cariological, allozyme and molecular) have been used. The phylogenetic trees based on different datasets partly contradict to each other, but certain clades are observed in all trees (see a review of Crespi and Fulton (2004)). These clades are as follows: (*S. confluentus*, *S. leucomaenis*), (*S. alpinus*, *S. malma*) and (*S. fontinalis*, *S. namaycush*). The last clade, most often, locates at the base of the *Salvelinus* tree. The topologies of the trees based on certain nuclear and mitochondrial genes data are also contradictory at a great degree. However, combined nuclear data yield a fully resolved phylogenetic tree for *Salvelinus.* This tree had the following topology: (*S. fontinalis*, *S. namaycush*), ((*S. confluentus*, *S. leucomaenis*), (*S. alpinus*, *S. malma*)). The authors (Crespi and Fulton, 2004) indicate that among the processes determining the contradictions between different genetic datasets, the historical hybridization events between the charrs of the genus *Salvelinus* represent an important factor.

It should be noted that in Crespi and Fulton work (as well as in many other studies conducted until the mid 2000s), not all main species of the charrs have been included in the analysis. In particular, *S. levanidovi* and *S. svetovidovi* are usually absent, and the *S. alpinus* – *S. malma* complex is represented by a single phylogenetic group of Arctic charr and Dolly Varden. In addition, the phylogenetic groups оf this complex are often different in various studies, and their evolutionary relationships remain obscured. Among recent investigations of phylogenetic relationships within the genus *Salvelinus*, a study on phylogenetic analysis of RAD-sequencing (RAD-seq) data in eight species including different phylogenetic groups of the *S. alpinus* – *S. malma* complex (Lecaudey et al. 2018) should be mentioned. All phylogenetic trees have been fully resolved. The clade (*S. levanidovi*, *S. leucomaenis*) is located near the root of the phylogenetic tree followed by the clade (*S. fontinalis*, *S. namaycush*), and then *S. confluentus* is disposed. Long-finned charr has a sister position regarding the clade of the *S. alpinus* – *S. malma* complex. The representatives of four phylogenetic groups of Arctic charr represent a single clade (*S. alpinus* complex), and southern Dolly Varden from Asia and northern Dolly Varden form the second clade (*S. malma* complex).

To minimize the negative impact of such factors as ‘taxon sampling’ (Wiens 2006; Heath et al. 2008) and ‘imperfect taxonomy’ (Funk and Omland 2003), all “good” charr species and all main representatives of the *S. alpinus* – *S. malma* complex (being confident of their phylogenetic and taxonomic identification) have been included in the analysis. The only exclusion is connected with the problem of taxonomic identification of a part of charr populations from the Kolyma basin. These populations bear CR haplotypes of mtDNA of northern Dolly Varden, which have been captured because of introgressive hybridization during the postglacial period. In particular, the charr population from Malyk Lake we preliminary refer to *S. a. erythrinus* (but not to *S. a. taranetzi*) despite ambiguous genetic markers data (Osinov et al. 2017, 2018, this study). Unfortunately, the samples of Arctic charr from Alaska and Arctic regions of Canada have not been included in our analysis. However, their phylogenetic similarity with the charrs of Taranets group from Asia is obvious (Osinov 2001; Brunner et al. 2001; Osinov et al. 2018).

The significant ILD test values indicated the absence of consistency between the topologies of the three (CR, ITS1, and RAG1) and two (ITS1 and RAG1, ITS1 and CR) gene trees. Phylogenetic analysis conducted in this study showed that the position of several taxa and clades in the gene trees, concatenated (ITS1+RAG1) trees and species trees was unstable, and support values (bootstraps or posterior probabilities), most often, were moderate or low.

Nevertheless, certain clades were defined in the majority of phylogenetic trees. In particular, the clade (*S. fontinalis*, *S. namaycush*) was revealed in the ITS1 gene trees, concatenated (RAG1+ ITS1) trees from PAUP and BEAST and species tree (RAG1, ITS1) of the *Salvelinus* from *BEAST analyses. The clade (*S. levanidovi*, *S. leucomaenis*) was present in the trees constructed based on nuclear genes data, and it had good support in several trees. In some trees based on nuclear (ITS1 and ITS1+RAG1) data, a sister taxon for the (*S. levanidovi*, *S. leucomaenis*) clade was *S. confluentus* with strong support. In general, the position of this taxon is contradictory. In some trees, it is close to *S. fontinalis* and *S. namaycush* (e.g. RAG1); and in CR tree, it is combined with southern Dolly Varden from North America into the East Pacific mtDNA clade. In the MDC-based species tree based on CR, RAG1 and ITS1 DNA fragments, *S. confluentus* is a sister taxon to the *S. alpinus* – *S. malma* complex. The data of this study, as well as the RAD-seq data (Lecaudey et al. 2018), indicate that the sister taxon to *S. leucomaenis* is *S. levanidovi* instead of *S. confluentus*. However, based on our data, the last taxon can represent a sister group to the two first taxa. It should be noted that the range of Levanidov’s charr is located inside of the white-spotted charr range, and the ranges of white-spotted charr and bull trout are located from both sides of the ancient Bering Land Bridge connected two continents from the Eocene to Late Miocene. The first appearance of the Bering Strait in the Late Miocene approximately 5.3 Mya (Gladenkov et al. 2002) facilitated the separation of the basal charr species distributed in Eurasia and North America.

Based on morphological analysis (Behnke, 1984), allozyme data (Crane et al. 1994; Osinov 2001) and nuclear RAG1 gene (Shedko et al. 2012), different forms of Dolly Varden (*S. malma* complex) and Arctic charr (*S. alpinus* complex) represent two presumably monophyletic groups. This conclusion is also supported by RAD-seq data (Lecaudey et al. 2018), but southern Dolly Varden from North America has not been included in the analysis (as well as in the RAG1 gene study). All three forms of Dolly Varden differ from various Arctic charr forms in certain meristic characters including smaller gill raker and pyloric caeca numbers. The phylogenetic similarity of southern Dolly Varden from Asia and North America is supported by the same diploid chromosome number (2n = 82) (Viktorovsky 1978; Cavender and Kimura 1989), and Behnke (1984) proposes a recent divergence of these forms. Nevertheless, according to Frolov (2000), karyological data support a polyphyletic origin of three Dolly Varden forms, and southern Dolly Varden from North America is phylogenetically similar to Taranets charr. According to ITS1 data, southern Dolly Varden from North America form a clade with Arctic charr and Taranets charr, and northern Dolly Varden form a monophyletic clade with southern Dolly Varden from Asia (Phillips et al. 1999; this paper).

A part of our data supported a monophyletic origin of the *S. alpinus* – *S. malma* complex, as well as monophyly of each of the subgroups (*S. alpinus* complex and *S. malma* complex). However, another part of our data contradicted with this hypothesis. A monophyletic origin of the supercomplex and two of its complexes was supported by only RAG1 data. A monophyletic origin of the *S. alpinus* – *S. malma* complex was also supported by combined RAG1+ ITS1 data, as well as by the species tree based on nuclear genes (but with the inclusion of long-finned charr). In addition, a monophyletic origin of the *S. alpinus* complex was defined in species tree from *Beast analysis based on the three genes. However, these clades had low posterior probability values in the species trees. The MDC-based species tree indicated a monophyletic origin of the three Dolly Varden forms (*S. malma* complex). The most precarious topological position was registered in southern Dolly Varden from North America, which could be transferred from the *S. malma* complex to the *S. alpinus* complex. It should be noted that *S. confluentus* and *S. svetovidovi* were located inside of the *S. alpinus* – *S. malma* complex clade in several trees.

The final comments of our data on the phylogeny of the charrs are as follows. The clade (*S. fontinalis*, *S. namaycush*) or the clade (*S. levanidovi*, *S. leucomaenis*) is located near the root of the *Salvelinus* tree. The former topology is according to the traditional opinion on the *Salvelinus* phylogeny (Crespi and Fulton, 2004), and the latter one is based on the RAD-seq data (Lecaudey et al. 2018) and on the part of the data of this study. Then *S. confluentus* or *S. svetovidovi* are located on the tree followed by the *S. alpinus* – *S. malma* complex clade. As it is mentioned above, the position of bull trout and southern Dolly Varden from North America is the most contradictory. The conflict between the topologies of different trees and low support values for several taxa and clades can be connected with different processes including incomplete lineage sorting and introgressive hybridization. These latter reasons are ignored by the classical methods of phylogenetic analysis, and the modern methods including the species tree inference from gene tree topologies lead to minimum incongruence due to ILS but do not take into account the consequences of introgressive hybridization (Leache et al. 2014; Yu et al. 2013) that cause possible incorrect topologies of the trees.

### Introgressive hybridization and phylogeny of charrs of the genus Salvelinus

The topology of the MDC-based species tree (number of reticulation nodes is equal to zero) differs from the topology of the phylogenetic tree obtained based on RAD-seq data (Lecaudey et al. 2018). The differences are connected with the position of the taxa located near the root of the *Salvelinus* tree, the order of joining of *S. confluentus* and *S. svetovidovi* and absence of monophyly of the *S. alpinus* complex in the MDC-based species tree. As is known, the application of the minimizing deep coalescence criterion for inferring species tree with five and more taxa is not statistically consistent (Than and Rosenberg 2011). A similar problem can appear during the use of the parsimony approach for the construction of phylogenetic networks. However, this approach is satisfactory in many cases (Yu et al. 2013). The phylogenetic networks obtained in this study differed in the topologies and sets of taxa participated in hybridization. Nevertheless, the number of possible topological shifts, as well as the number of taxa entered into introgressive hybridization was restricted. According to networks with different number (from one to three) of reticulation nodes, four taxa participated in hybridization most often. They were as follows: southern Dolly Varden from North America, bull trout, northern Dolly Varden and the charrs of the Taranets group, i.e. the taxa with the most frequently occurred differences in their topological position in the phylogenetic and species trees.

In the MDC-based species tree and phylogenetic networks, *S. confluentus* located in two main positions. According to the first position, it is a sister taxon to the *S. alpinus* – *S. malma* complex. The second position indicates that it is a sister taxon to the Taranets charr clade including (in this study) Taranets charr of Chukotka and two species from Lake El’gygytgyn. Based on the majority of networks, *S. confluentus* entered into hybridization with southern Dolly Varden from North America, with a representative of Taranets charr group (one case) and with a common ancestor of *S. levanidovi* and *S. leucomaenis* (one case). The nuclear DNA data (Crespi and Fulton, 2004; Lecaudey et al. 2018; this study) and allozyme data (Crane et al. 1994) indicate that bull trout belongs to the basal species group, and it is not referred to the *S. alpinus* – *S. malma* complex. However, this conclusion contradicts with the mtDNA data: the bull trout forms a clade (East Pacific (EP)) together with southern Dolly Varden from North America, and this clade is a sister group to the Arctic clade combining the haplotypes of the Taranets charr group. The mito-nuclear discordance can be eliminated based on the assumption (Phillips et al. 1995) that mtDNA of the East Pacific haplogroup is not the native mtDNA of the bull trout. This mtDNA, most likely, was captured by bull trout during the hybridization with southern Dolly Varden from North America (that is supported in the majority of the networks) or was obtained directly from Taranets charr. However, it remains unclear whether mtDNA of the EP haplogroup is native to the southern Dolly Varden from North America or it was transferred from Taranets charr in the past.

It should be noted that a sister-group relationship between the East Pacific and Arctic clades rejects monophyly of *S. alpinus* complex and *S. malma* complex. The level of genetic divergence between EP and Arctic haplogroups is high (p-distance = 2%, on average). Therefore, the transfer of mtDNA from Taranets charr to southern Dolly Varden from North America (or to bull trout and then to southern Dolly Varden) or in the opposite direction occurred more than 1 Mya, and it was not associated with the last glacial period. According to some authors, the Bering group haplotypes represent original mtDNA of southern Dolly Varden from North America, and the EP haplotypes were transferred from bull trout to this form in the glacial refugium (Redenbach and Taylor 2002). This hypothesis can be reliable, but it means that introgression of mtDNA during hybridization between southern Dolly Varden from North America and bull trout could occur repeatedly. It also remains unclear whether mtDNA of the Bering group is native to southern Dolly Varden from North America. In particular, at least a part of the BER haplotypes of this form, most likely, was obtained because of hybridization with northern Dolly Varden in postglacial time (e.g., Taylor and May-McNally 2015). Two BER haplotypes revealed in the population of southern Dolly Varden from Graham Island have a basal position at the Bering clade, and their occurrence before the last glacial period cannot be excluded. According to many researchers, a glacial refugium located in this region (e.g. Shafer et al. 2010).

In the MDC-based species tree, all forms of Dolly Varden produced a single clade supporting their monophyletic origin. Two lineages of southern North America Dolly Varden (the first lineage with Bering mtDNA haplotypes, and the second lineage with East Pacific mtDNA haplotypes) are joined into a single clade in the species tree and the most of the networks. Although an assumption on the origin of all forms of Dolly Varden from a common ancestor seems the most probable, the origin of southern Dolly Varden from North America remains uncertain. Among three genes (DNA fragments) included in our analysis, only nuclear RAG1 gene data supported the monophyly of the *S. malma* complex. Based on the majority of networks, hybridization between southern Dolly Varden from North America and northern Dolly Varden, as well as with bull trout or with Taranets charr, took place. Other genetic data support hybridization between three first taxa. In particular, a zone of admixture between southern and northern Dolly Varden is found in the region in and around the Gulf of Alaska (Taylor and May-McNally 2015), and an intergradation zone between southern Dolly Varden and bull trout is described in British Columbia (Redenbach and Taylor 2002).

In addition, our data indirectly indicated that southern Dolly Varden from North America was represented by two glacial lineages originated from different refugia and, most likely, reproductively isolated from each other. In this study, two small samples of southern Dolly Varden from North America were presented by Canadian colleagues and were collected in two adjacent streams of Graham Island. The distance between their mouths was approximately 5 km, and apparent barriers for mutual migrations and gene exchange between populations of southern Dolly Varden were not observed (Dr. A.Costello, personal communication). The information about a possible occurrence of anadromous Dolly Varden form (at least in one of the two streams) participated in the present gene flow is absent. Each of the populations reveales the haplotypes of only one haplogroup (Bering or East Pacific). Other populations of southern Dolly Varden (e.g., from Vancouver Island) include mtDNA of only one haplogroup, although some of them have haplotypes of both haplogroups (Redenbach and Taylor 2002), indicating the gene flow in the past. This pattern of genetic differentiation can be explained by gene drift or by the presence of two partly reproductively isolated forms of Dolly Varden in this region. The presence of two “ mtDNA lineages” of southern Dolly Varden in North America is supported by microsatellite DNA analysis (Taylor and May-McNally 2015).

In the majority of phylogenetic trees and networks, northern Dolly Varden joins with southern Dolly Varden from Asia. The trees constructed based on mtDNA (or including mtDNA data) represent the exclusion: the Bering haplogroup of northern Dolly Varden has a sister position to the clade including the haplotypes of Arctic charr from Eurasia (Atlantic and Siberia haplogroups). Based on the apparent contradiction between the allozyme (Osinov 2001) and mtDNA data (Brunner et al., 2001), an assumption about the capture of mtDNA of Arctic charr by northern Dolly Varden in the past has been proposed (Shedko et al. 2007; Osinov et al. 2015). In the MDC-based species tree and the majority of phylogenetic networks, the clade of Arctic charr from Europe and Siberia is a sister group to the clade composed of three (or two) Dolly Varden forms. This topology, most likely, does not indicate a close phylogenetic relationship between these taxa: the capture of mtDNA of Arctic charr from Eurasia by northern Dolly Varden could occur in the past. Only one network supports this assumption. According to molecular clock data, the capture of ‘alien’ mtDNA by northern Dolly Varden occurred 1.2–1.9 Mya (Osinov et al. 2015). During this period, the divergence of Arctic charr from Siberia, Europe and populations from Atlantic coast of North America (haplogroups Siberia, Atlantic and Acadia), as well as the divergence of the Arctic and East Pacific haplogroups, began. According to numerous data, introgression of mtDNA in opposite direction (from northern Dolly Varden to Arctic charr) occurred in the postglacial time due to secondary contacts and hybridization between northern Dolly Varden and different forms of Arctic charr (Taylor et al. 2008; Alekseyev et al. 2009; Osinov et al. 2017, 2018; Esin et al. 2017).

All networks supported sister relationships between northern Dolly Varden and southern Dolly Varden from Asia, but none of them suggested hybridization events between these forms. To simplify the analysis (at a preliminary stage), we excluded an individual of southern Dolly Varden with mtDNA haplotype of the northern Dolly Varden because of an apparent hybridization event. It should be noted that mtDNA haplotypes of northern Dolly Varden are revealed in many populations of southern Dolly Varden from the Kuril Islands and Sakhalin, and, in some cases, they have high frequencies (Shedko et al. 2007; Osinov and Mugue, 2008). An occurrence of introgressive hybridization between southern Dolly Varden and northern Dolly Varden from Asia is supported by morphological analysis (Pichugin et al. 2008) and allozyme data (Osinov 2001). The present ranges of these forms can be partly overlapped in Primorye, Sakhalin and the northern Kuril Islands, while the populations might have been strongly reproductively isolated. Based on the opinion of Safronov and Zvezdov (2005), three charr species (southern Dolly Varden (*S. krascheninnikovi*), its resident form (*S. curilus*) and a new species, Sakhalin charr *S. vasiljevae*, described by the authors) are jointly distributed in certain rivers of northwestern Sakhalin. In our opinion, the existence of different forms in several Sakhalin rivers, most likely, can be explained by the occurrence of a zone of the secondary contact between northern Dolly Varden and southern Dolly Varden (Osinov and Mugue 2008; Pichugin et al. 2008). An occurrence of the three or only two charr (Dolly Varden) forms in Sakhalin, as well as a degree of their reproductive isolation, will be specified only after the population genetic analysis.

The long-finned charr has different topological positions in the nuclear RAG1 gene tree and mtDNA control region tree. Two networks indicate a hybridization event between *S. svetovidovi* and Arctic charr from Siberia or the ancestor of Taranets charr. As it is mentioned above, the AC1 haplotype of the nuclear RAG1 gene (which is the main haplotype in all representatives of the *S. alpinus* complex) is fixed in long-finned charr and sympatric small-mouth charr *S. elgyticus*. A presence of this haplotype in *S. svetovidovi* can be connected with ILS, but, most likely, it is a consequence of introgressive hybridization with the small-mouth charr. Based on the data on mtDNA and microsatellites, at present, hybridization between three charr species of Lake El’gygytgyn is absent, but hybridization and introgression of several genes, most likely, was possible during the postglacial period when these species entered into secondary contacts (Osinov et al. 2015). A network indicates an occurrence of hybridization between long-finned charr and the common ancestor for northern Dolly Varden and southern Dolly Varden from Asia.

During the analysis of phylogeny of charrs based on RAD-seq data, the four-taxa D-statistics test (Durand et al. 2011) has been applied to analyze a level of gene introgression between different taxa (Lecaudey et al. 2018). This level is not high ranging from 1.66% between *S. namaycush* and *S. leucomaenis* to 4.22% between *S. alpinus* (ARC) (i.e., Taranets charr) and *S. malma* (BER) (i.e., northern Dolly Varden). In this study, gene introgression between two former species, as well as between several other species (e.g., between *S. svetovidovi* and *S. namaycush* (2.19%) or between *S. svetovidovi* and *S. levanidovi* (2.56%)), was not revealed. In both studies, gene introgression between northern Dolly Varden and different phylogenetic groups of Arctic charr (*Salvelinus alpinus* complex) was demonstrated. Thus, the data of the two analyzes partially coincided, but partially were different. Among possible reasons of the disagreement, two of them should be noted. At first, southern Dolly Varden from North America (a taxon with the most various and intensive hybridization relationships according to our data) has not been included in the RAD-seq analysis. In addition, two other species from Lake El’gygytgyn (hybridization between long-finned charr and these species seems more probable than hybridization between *S. svetovidovi* and a species from North America) also have not been included. Secondly, the RAD-seq data can include the data on a small number of short mtDNA fragments, but this possibility is real not for all taxa. In some taxa, mtDNA does not have a restriction site for *Sbf1* endonuclease used for the analysis. In this study, mito-nuclear discordances played an important role in the analysis of hybridization events between different taxa. In general, both studies, as well as other data, indicate that many charr species and representatives of all phylogenetic groups of the *S. alpinus* – *S. malma* species complex with present overlapping or at least adjoining ranges entered into secondary contacts with subsequent hybridization and gene introgression. Because of the contradiction in the results of this and RAD-seq studies, the levels of gene introgression between different taxa should be specified. The consequences of the hybridization can be variable leading to both extinction (Todesco et al. 2016) and speciation and adaptive radiation (Abbott et al. 2013; Seehausen 2013). Gene introgression is at least partly responsible for the contradictions in the topologies of different gene trees and for the troubles, which appear during the analysis of phylogeny of charrs of the genus *Salvelinus*.

## Supporting information

Online Resourse 1

## ACKNOWLEDGEMENTS

We thank S.S. Alekseyev, M.Yu. Pichugin, O.A. Radchenko, P.V. Grigor’ev, L. Bernatchez, A. Costello, E.B. Taylor, U.K. Schliewen, R. Reiter and S. Yamamoto for help in the collection of the material. The research was supported by the Russian Foundation for Basic Research (project no. 17-04-00063).

## SUPPORTING INFORMATION

### Online Resource 1

Figure 1. Approximate modern ranges of the representatives of two species complexes. (A) The Arctic char complex: the Arctic charr of Eurasia, including *S. a. alpinus* from Europe (“Atlantic” mtDNA haplogroup) and *S. a. erythrynus* from Siberia (“Siberia” haplogroup), Taranets charr, *S. a taranetzi* (Arctic haplogroup) and the Arctic charr from New England and southeastern Canada, *S. a. oquassa* (Acadia haplogroup). (B) The Dolly Varden complex: the southern Dolly Varden of North America, *S. m. lordi* (Bering and East Pacific haplogroups), the northern Dolly Varden, *S. m. malma* (Bering haplogroup), the southern Dolly Varden of Asia, *S. m. krascheninnikovi* (West Pacific haplogroup).

### Online Resource 2

Figure 2. (A) MDC-based species tree and phylogenetic networks based on three genes with (B) one and (C) two reticulation nodes. Rich Newick format includes reticulation nodes (#H) and inheritance probabilities (in bold). Abbreviations: a, *S. fontinalis*; b, *S. namaycush*; c, *S. levanidovi*; d, *S. leucomaenis*; e, *S. svetovidovi*; f, *S. malma krascheninnikovi*; g, *S. malma malma*; i, *S. confluentus*; j, *S. malma lordi* (EP); k, *S. malma lordi* (BER); l, *S. boganidae*; m, *S. elgyticus*; n, *S. alpinus taranetzi*; p, *S. alpinus alpinus*; h*, S. alpinus erythrinus*.

### Online Resource 3

Figure 3. Phylogenetic networks based on three genes with three reticulation nodes. Rich Newick format includes reticulation nodes (#H) and inheritance probabilities (in bold). Abbreviations: a, *S. fontinalis*; b, *S. namaycush*; c, *S. levanidovi*; d, *S. leucomaenis*; e, *S. svetovidovi*; f, *S. malma krascheninnikovi*; g, *S. malma malma*; i, *S. confluentus*; j, *S. malma lordi* (EP); k, *S. malma lordi* (BER); l, *S. boganidae*; m, *S. elgyticus*; n, *S. alpinus taranetzi*; p, *S. alpinus alpinus*; h*, S. alpinus erythrinus*.

