## Supplementary material for "Charrs of the genus *Salvelinus* (Salmonidae): hybridization, phylogeny and evolution": Online Resourse 1

A

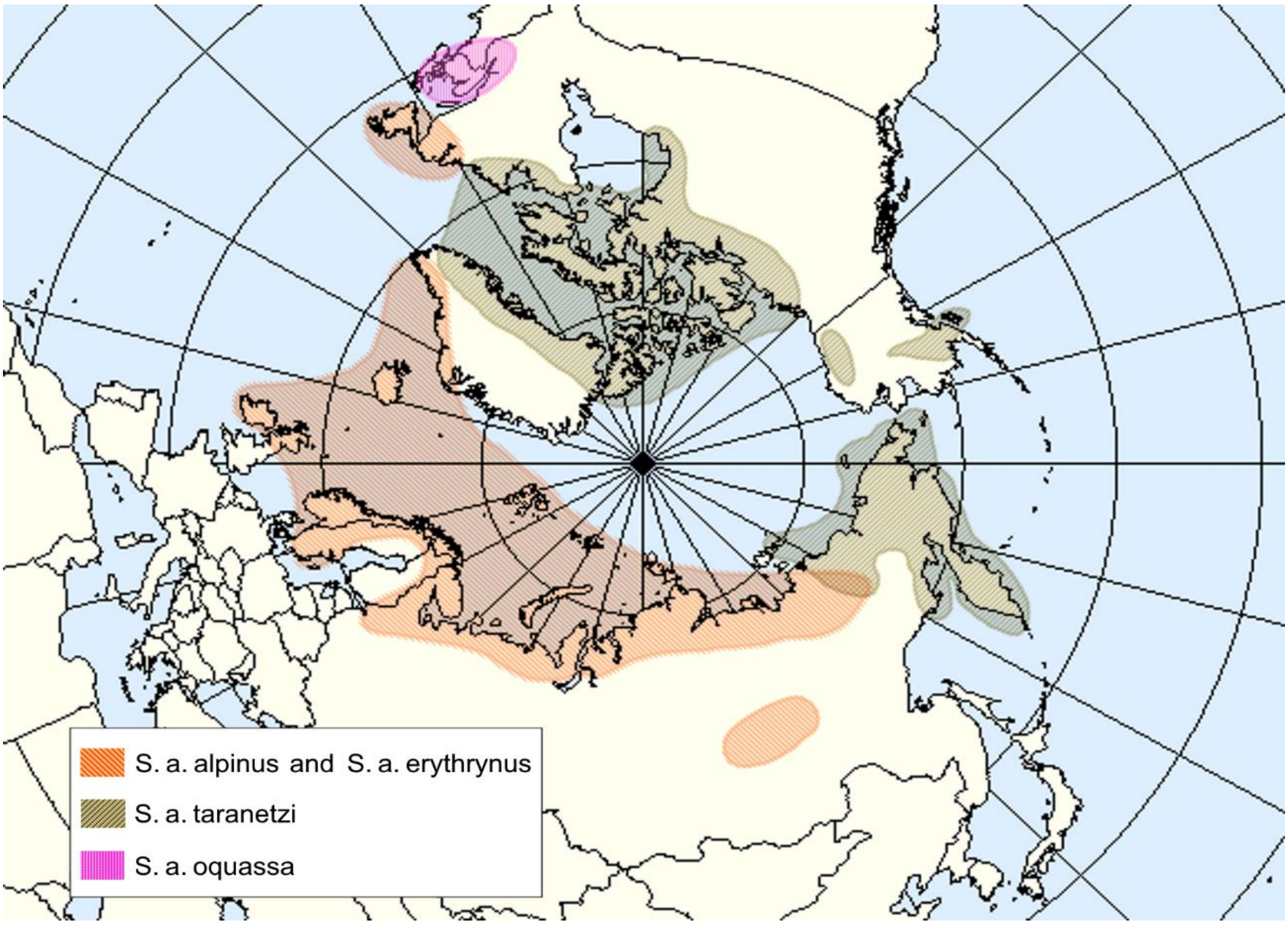
S. a. alpinus and S. a. erythrynus
S. a. taranetzi
S. a. oquassa

B

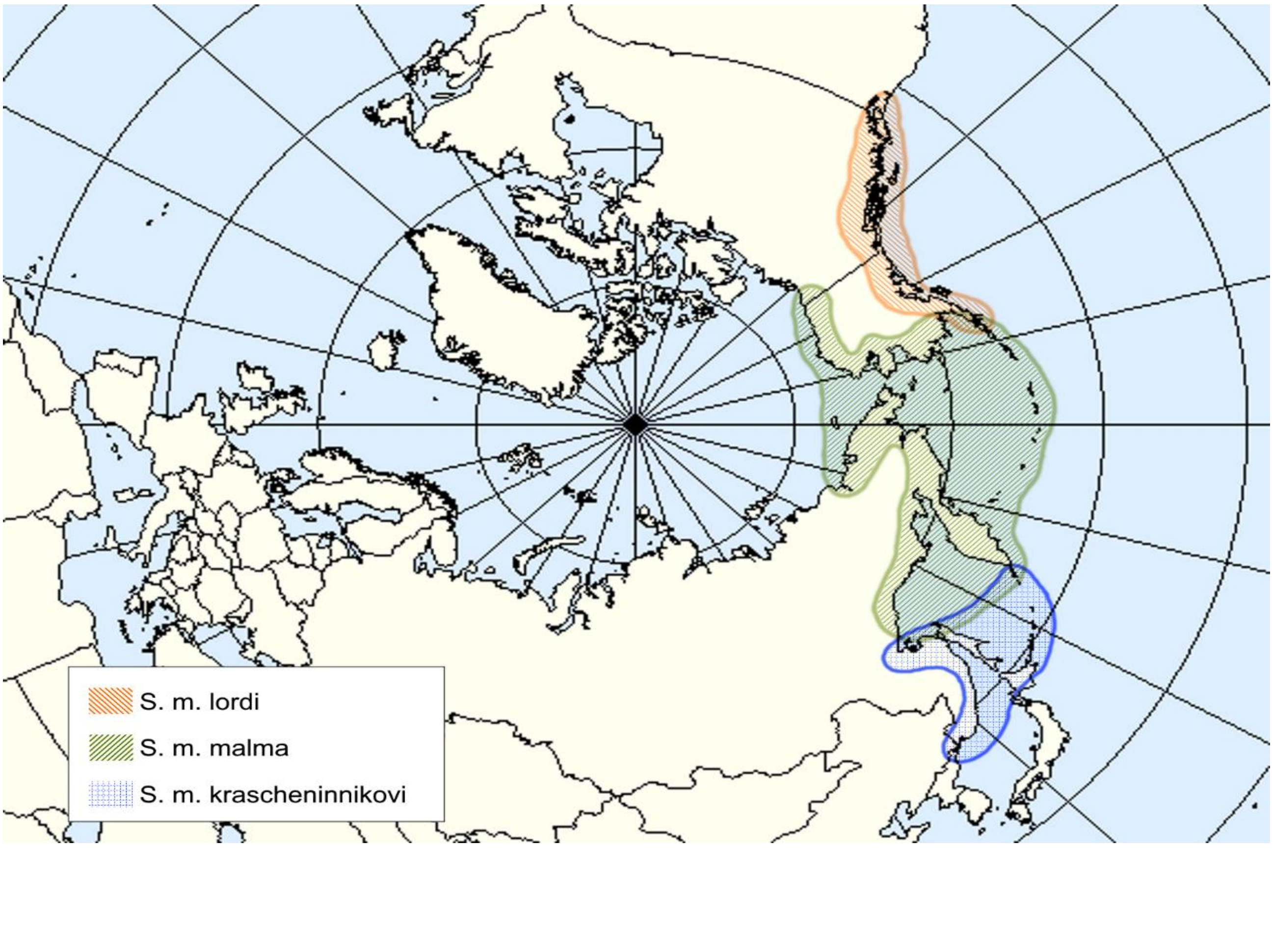
S. m. lordi
S. m. malma
S. m. krascheninnikovi

**Figure S1.** Approximate modern ranges of the representatives of two species complexes. (A) The Arctic char complex: the Arctic charr of Eurasia, including *S. a. alpinus* from Europe (Atlantic mtDNA haplogroup) and *S. a. erythrynus* from Siberia (Siberia haplogroup)**,** Taranets charr, *S. a taranetzi* (Arctic haplogroup) and the Arctic charr from New England and southeastern Canada, *S. a. oquassa* (Acadia haplogroup). (B) The Dolly Varden complex: the southern Dolly Varden of North America, *S. m. lordi* (Bering and East Pacific haplogrups)**,** the northern Dolly Varden, *S. m. malma* (Bering haplogroup)**,** the southern Dolly Varden of Asia, *S. m. krascheninnikovi* (West Pacific haplogrup)**.**
